# Brain and neuronal expression and localization of de-S-acylating enzymes

**DOI:** 10.64898/2026.05.31.729046

**Authors:** Gabriella Santander Herrera, Nisandi N. Herath, Amelia H. Doerksen, Simone I.M. Clarke, Yasmeen Alshehabi, Milena Rabu, Julia E. Fux, Charlotte A. Townsend Bennie, Dale D.O. Martin, Shaun S. Sanders

## Abstract

S-acylation is a reversible posttranslational lipid modification important in the nervous system that dynamically regulates protein localization and function. Aberrant S-acylation has been implicated in several neurological conditions. While several de-S-acylases (deacylases hereafter) have been identified, little is known regarding their expression and localization in the brain and in neurons. Here, we characterized the expression, localization, and S-acylation of cytosolic deacylases, including acyl-protein thioesterases APT1, APT2, and APT1L and α/β hydrolase domain-containing proteins ABHD7, ABHD10, ABHD13, ABHD16A, and ABHD17A-C. Mouse brain RNA sequencing data reveal high expression of *Lypla1*/APT1, *Lypla2*/APT2, *Ephx4*/ABHD7, *Abhd16a*, and *Abhd17A-C* in the brain, whereas *Lyplal1*/APT1L, *Abhd10*, and *Abhd13* were expressed at very low levels. At the protein level, APT1 and ABHD16A levels are highest in the cerebellum with ABHD17A levels lowest in this region while APT2 levels are highest in the hippocampus. However, all four are abundant in cultured hippocampal neurons. Deacylases are localized throughout neurons on punctate structures, with APT2 and ABHD17C localized to the Golgi. Finally, all ten cytosolic deacylases are themselves S-acylated. These data characterizing deacylase expression, localization, and S-acylation in neural contexts, provides a foundation for future studies investigating deacylase neuronal functions and potential roles in neurological disease.

## Introduction

Neurons are large, complex cells that require efficient and precise protein trafficking and localization in response to changes in neuronal activity. Dynamic posttranslational modifications are key mechanisms governing this process. S-acylation, commonly S-palmitoylation, is a reversible lipid modification where long-chain fatty acids are added to cysteine residues to increase protein hydrophobicity. This, in turn, dynamically regulates protein membrane association and sorting to membrane microdomains, protein-protein interactions, protein stability, and protein function [1–4]. S-acylation plays a key role in the nervous system, where ∼50% of synaptic proteins, ∼80% of axon initial segment proteins, and ∼65% of neuronal trafficking machinery are likely S-acylated [2,3]. Furthermore, aberrant S-acylation has been implicated in a range of neurological disorders, including X-linked intellectual disability, psychiatric disorders, and neurodegenerative diseases [1–5].

S-acylation is mediated by S-acylating enzymes and de-S-acylating enzymes (hereafter deacylases). There are three classes of deacylases: acyl-protein thioesterases (APTs), α/β hydrolase domain-containing proteins (ABHDs), and palmitoyl protein thioesterases (PPTs) [1,6]. PPT1 is lysosomal and deacylates proteins prior to degradation [6]. APT1/LYPLA1, APT2/LYPLA2, APT1L/LYPLAL1, ABHD7/EPHX4, ABHD10, ABHD13, ABHD16A/BAT5, and ABHD17A-C are the cytosolic deacylases [1,7–9]. APT1 and APT2 are well characterized with many known substrates, including the GTPase Ras and HTT, the protein mutated in Huntington disease [6,10]. ABHD17A-C also have many known substrates, including the excitatory postsynaptic scaffold PSD95 [10,11]. APT1L deacylates the large conductance calcium-activated potassium channel [12] and ABHD13 and ABHD16A deacylate the SNARE protein SNAP25 [7]. ABHD16A also deacylates interferon-inducible transmembrane proteins [8], ABHD7 deacylates Lamin A [9], and mitochondrial ABHD10 deacylates peroxiredoxin-5 [13].

Several deacylases have been implicated in neurological disorders [6]. For example, APT1 and APT2 are therapeutic targets to restore mutant HTT S-acylation in Huntington disease [5,14,15] and mutations in *ABHD16A* cause spastic paraplegia [16], while *ABHD13* and *ABHD17B* are differentially expressed in amyotrophic lateral sclerosis [17]. However, little is known regarding the expression and localization of these enzymes in the brain and neurons, limiting understanding of their role in this context. Thus, we sought to characterize the expression and localization of the deacylases across the brain and in neurons.

## Methods

Plasmid, antibody, and reagent information as well as buffer and media compositions and primer and gRNA sequences are in the key resources table in the supplementary material.

### RNA sequencing expression analysis

Mouse brain region bulk RNA sequencing (RNAseq) data were downloaded from proteinatlas.org (rna_mouse_brain_hpa.tsv) [18] as in BrainPalmSeq [19]. Protein-coding transcripts per million (pTPM) for *Lypla1, Lypla2, Lyplal1, Ephx4, Abhd10, Abdh13, Abhd16a, Abhd17a, Abhd17b*, and *Abhd17c* were plotted on a heat map.

### Animals

Animal use was approved by the University of Guelph Animal Care Committee according to the Canadian Council on Animal Care guidelines. Brain tissues from 6–8-month-old male and female C57BL/6 mice were dissected and flash frozen. Female timed pregnant Sprague-Dawley rats were purchased (Charles River Laboratories, OH, USA) and hippocampal tissue was collected from day 18 embryos (E18) for primary cultures.

### Tissue lysis

Brain tissues were homogenized in ice-cold homogenization buffer with 4% Roche cOmplete EDTA (ethylenediaminetetraacetic acid)-free protease inhibitor cocktail (PIC) before adding sodium dodecyl sulfate (SDS) to a final concentration of 2.5% to solubilize proteins. DNA was sheared by sonicating at 25% power until clear. Protein concentrations were determined using the *DC* protein assay according to the manufacturer’s instructions. 30 μg of protein was diluted to 100 μL and denatured in Laemmli Sample Loading Buffer with 1% β-mercaptoethanol at 95°C for 5 minutes for SDS-PAGE (polyacrylamide gel electrophoresis) and immunoblotting.

### Molecular biology and plasmids

N-terminally HA (hemagglutinin)-tagged cDNA plasmids with an EF1ɑ(Elongation Factor 1ɑ) promoter were gifts from Dr. Jennifer Greaves (Coventry University) and the mCherry plasmid (pFEmCherryW; EF1ɑ) was a gift from Dr. Gareth Thomas (Temple University). N-terminally HA-tagged *ABHD7* and *ABHD17B* plasmids were generated by PCR subcloning into the BamHI site of pEF-BOS. C-terminally HA-tagged *LYPLA1, LYPLA2*, and *ABHD10* cDNA plasmids were generated by PCR subcloning from pEF-BOS into the XhoI/NotI sites of FEW-HA (EF1ɑ) to avoid interrupting N-terminal S-acylation and/or mitochondrial targeting. LentiCas9-Blast was a gift from Dr. Feng Zhang and plasmids with dual CRISPR/Cas9 guide RNAs (gRNA; pCLIP-DUAL-sgRNA) targeting human *LYPLA1, LYPLA2, ABHD16A, ABDH17A*, or *eGFP* (non-targeting [NT], negative control) were purchased from the University of Ottawa GEM Facility. All constructs were verified by Sanger or nanopore sequencing.

### Hippocampal neuron culture and transfection

Hippocampal neurons were cultured from E18 rat hippocampi in complete Neurobasal medium (NBM) at 37°C in 5% CO_2_ and treated with the mitotic inhibitor 5-Fluoro-2’-deoxyuridine on day *in vitro* (DIV) 5 [20]. Neurons were lysed on DIV5, 10, 15, and 20 in 2% SDS lysis buffer with 4% PIC (2SB) and prepared for SDS-PAGE and immunoblotting as above. For immunocytochemistry, neurons were plated on poly-L-lysine coated glass coverslips and transfected on DIV13 using Lipofectamine 2000 Transfection Reagent (L2K). 750 ng of deacylase and 500 ng pFEmCherryW DNA were diluted in 50 μL NBM and mixed dropwise with an equal volume NBM containing 3 μL L2K. DNA-L2K complexes were incubated for 15 minutes at 37°C before adding dropwise to neurons. Media was replaced 90 minutes later with a 1:1 mixture of conditioned and fresh complete NBM. Neurons were fixed 24 hours later in parafix for 10 minutes at 21°C for immunocytochemistry.

### Cell surface biotinylation

Neurons were washed three times with ice-cold artificial cerebral spinal fluid (aCSF) prior to biotinylation of surface exposed proteins using 0.75 mg/mL Sulfo-NHS-SS-Biotin in ice-cold aCSF for 30 minutes. Neurons were washed once with ice-cold aCSF with 50 mM tris-HCl pH 7.0 and twice with ice-cold aCSF without tris, lysed in 2SB, and prepared for SDS-PAGE and immunoblotting as above. 300 μg of protein was diluted 1:20 in dilution buffer and biotinylated proteins purified using NeutrAvidin beads for 3 hours at 4°C. Beads were washed three times with dilution buffer containing 0.5 M NaCl, once without NaCl, and surface proteins eluted in elution buffer at 37°C for 10 minutes and denatured as above for SDS-PAGE and immunoblotting.

### HEK293T and HAP1 cell culture and transfection

HEK293T (human embryonic kidney 293T) cells were cultured in complete DMEM (Dulbecco’s modified eagle medium) and HAP1 (human haploid) cells in complete IMDM (Iscove’s Modified Dulbecco’s medium) at 37°C in 5% CO_2_. HEK293T cells were transfected using CaPO_4_ [20]. 3 μg of deacylase DNA was diluted in 250 μL 244 mM CaCl_2_ and combined dropwise with 250 μL 2x HEPES-Buffered Saline with mixing before adding dropwise to 80-90% confluent cells seeded the previous day. The media was replaced after 4 hours with complete DMEM. Bioorthogonal labeling was performed the following day. HAP1 cells were transfected using PEI (polyethyleneimine). A PEI-DNA-IMDM mixture with a 1:5 ratio of PEI:DNA was made by diluting PEI to 1 mg/mL in media, adding DNA, vortexing, and incubating for 10 minutes at 21°C. The PEI-DNA complexes were added dropwise to the cells and 4 hours later the media was replaced with fresh complete IMDM.

### Gene knockout in HEK293T or HAP1 cells

*LYPLA1* knockout HEK293T cells and *LYPLA2* or *ABHD16A* knockout HAP1 cells were generated as previously described [21]. 62,000 cells per well of a 6-well plate were transfected with 1.2 μg of Lenti-Cas9-BLAST and 0.8 μg of pCLIP-DUAL-sgRNA. Cells were selected 48 hours later with 2 μg/mL puromycin until all cells in an untransfected well died. Cells were then maintained as a polyclonal population with a matched NT control line. Cells were lysed in 2SB and prepared for SDS-PAGE and immunoblotting as above. For immunocytochemistry, NT and *ABHD16A* or *LYPLA2* knockout HAP1 cells were plated on poly-L-lysine coated glass coverslips and fixed 24 hours later, as above, for immunocytochemistry.

### TIDE (Tracking of Indels by Decomposition) analysis

NT or knockout HEK293T or HAP1 cells were lysed in genomic DNA buffer and protein was digested with 100 μg/mL Proteinase K for 1 hour at 21°C before inactivating for 10 minutes at 95°C. 3 volumes of ice-cold 100% ethanol was added to precipitate genomic DNA overnight at - 20°C. DNA was pelleted at 4°C for 30 minutes at 13,000 rpm and washed twice with 75% ethanol before dissolving in water. Primers amplifying each gRNA-targeted region in the *LYPLA1, LYPLA2*, and *ABHD16A* genes were used in PCR with Q5 DNA Polymerase. Amplified DNA was purified by PCR Cleanup, Sanger sequenced, and the indel mutation rate relative to NT control was determined using TIDE [22].

### Immunocytochemistry and imaging

Fixed neurons or HAP1 cells were permeabilized with 0.25% Triton X-100 in phosphate buffered saline (PBS) for 10 minutes at 21°C, blocked in 2% gelatin in PBS for 1 hour at 21°C, and incubated overnight at 4°C in the indicated primary antibodies diluted in blocking buffer. Coverslips were subsequently washed and incubated with secondary antibodies diluted in blocking buffer for 1 hour at 21°C before washing and mounting onto glass microscope slides using PermaFluor Mounting Medium. HAP1 cells were imaged using a wide-field Nikon ECLIPSE T*i*2 fluorescent microscope with a 60x oil immersion objective (1.4 numerical aperture [NA], PL-APO) and neurons were imaged using 0.4 μm spaced Z-stacks with a Leica Microsystems STELLARIS 5 confocal microscope with LIGHTNING using a 63x oil immersion objective (1.4 NA, HC PL-APO). All neuron images are maximum intensity projections of LIGHTNING images modified for brightness and contrast using ImageJ Fiji [23] and are representative of 5-7 neurons.

### Bioorthogonal labeling and click chemistry

Transfected HEK293T cells were subjected to bioorthogonal labeling as previously described [24]. Cells were lipid depleted (DMEM with 5% charcoal stripped FBS) for 30 minutes before labeling with 100 μM saponified alkynyl-palmitate, alkynyl-stearate, or non-alkyne palmitate (negative control) bound to fatty acid free bovine serum albumin (BSA) for 4 hours. Cells were lysed in EDTA-free 2SB and prepared for SDS-PAGE and immunoblotting as above. 200-500 μg protein was reacted with 100 μM biotin azide plus using 1 mM CuSO_4_, 1 mM sodium L-ascorbate, and 0.1 mM of TBTA (tris(benzyltriazolylmethyl)amine) for 1 hour at 21°C before stopping with 10 mM EDTA. Protein was precipitated by adding ice cold acetone to 80%, protein pellets were dissolved in 2SB, and biotinylated proteins purified as above. S-acylated proteins were eluted in 40 μL hydroxylamine elution buffer with 4% PIC for 1 hour at 21°C and denatured as above for SDS-PAGE and immunoblotting.

### SDS-PAGE and immunoblot

Samples subjected to SDS-PAGE were transferred to nitrocellulose membranes. Membranes were stained with Ponceau for total protein, blocked in 5% skim milk in tris buffered saline (TBS) with 0.05% tween-20, and immunoblotted overnight at 4°C with indicated primary antibodies diluted in 1% BSA/TBS. Membranes were washed and probed with the appropriate secondary antibodies diluted in blocking buffer. Blots were imaged using Clarity Western ECL substrate with a ChemiDoc XRS+ with Image Lab Software and quantified using Image Studio Software.

### Statistical analysis

All data were graphed and analyzed using GraphPad Prism. Mean with standard deviation was plotted and statistical tests used are indicated in the figure legends. Cells from separate passages, individual animals, or independent neuronal cultures were considered biological replicates.

## Results & Discussion

We first compared deacylase mRNA expression across the mouse brain using bulk RNAseq data [18] (Figure 1). *Lyplal1, Abhd10*, and *Abhd13* are expressed at low levels across the brain with *Abhd17c* expressed at moderate levels in a subset of brain regions. *Lypla1, Lypla2, Abhd16a, Abhd17a*, and *Abhd17b* are well expressed across all regions, whereas *Ephx4* is expressed at high levels in the amygdala, basal ganglia, cerebral cortex, corpus callosum, and hippocampus but at low levels in other tissues. Comparing across genes, *Abhd17a* is the highest expressed across the brain, particularly within the cerebrum, and *Abhd16a* expression in the cerebellum is the highest of all genes across all regions. This suggests that APT1L, ABHD10, and ABHD13 may not play a prominent role in brain protein deacylation. In contrast, APT1, APT2, ABHD7, ABHD16A, and ABHD17A-C may represent the major brain deacylating enzymes, with individual enzymes potentially particularly important in specific regions, i.e., ABHD16A in the cerebellum.

**Figure 1.**
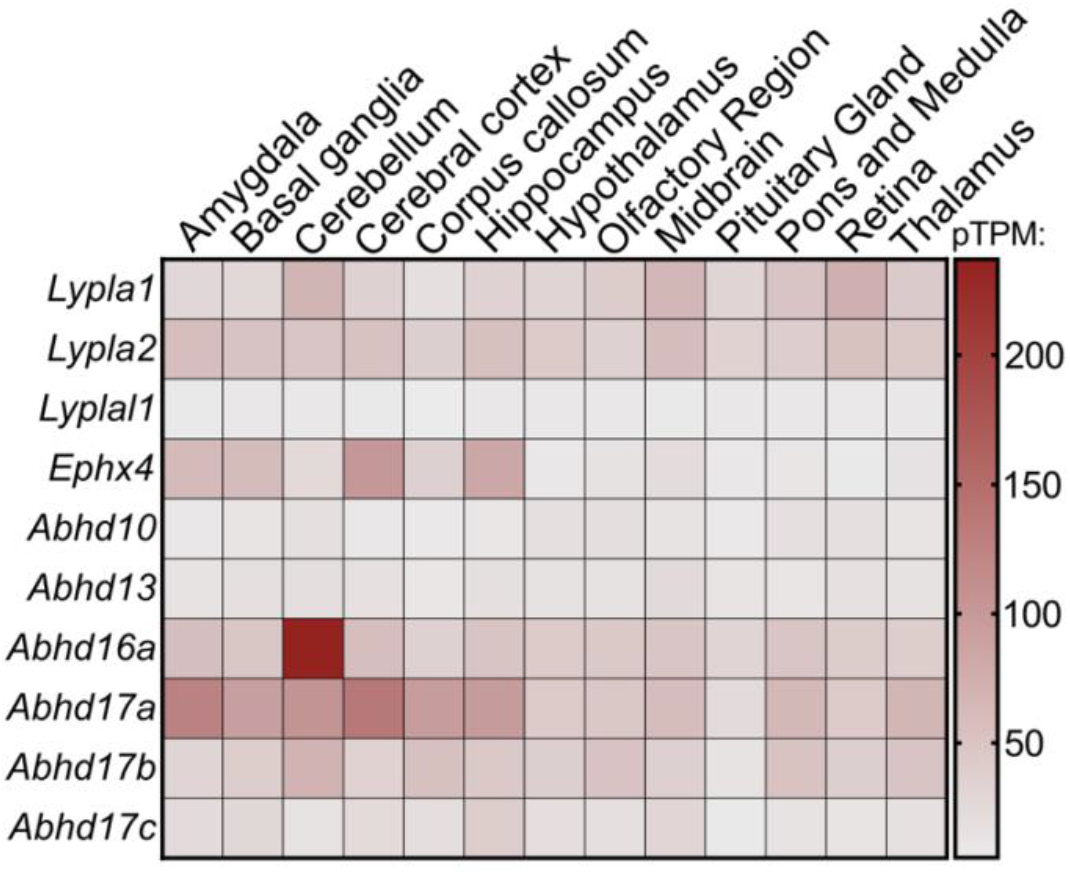
Deacylase RNA expression in mouse brain. Mouse bulk RNAseq pTPM data for the deacylases were compiled from the Sjöstedt ProteinAtlas [18] and plotted on a heat map.

We next sought to confirm the RNAseq data at the protein level. First, we validated commercial deacylases antibodies for immunoblotting using lysates from mock and HA-deacylase transfected HEK293T cells and hippocampal neurons. The APT1, APT2, ABHD7, ABHD13, ABHD16A, and ABHD17A antibodies recognized the corresponding HA-tagged protein whereas the ABHD10, ABHD17C, and the second ABHD13 and ABHD17A antibodies did not (Fig. S1A-J). The ABHD7 and ABHD13 blots had many non-specific bands hindering identification of the endogenous band (Fig. S1C & F). Further validation showed that the primary band at the predicted molecular weights for the APT1, APT2, and ABHD16A antibodies was reduced in the corresponding polyclonal knockout cells, indicating on-target binding (Fig. S1K-N; indel efficiency in Table S1). We also tested the ABHD16A and APT2 antibodies for immunocytochemistry in knockout cells but both exhibited non-specific staining (Fig. S2).

We next assessed protein levels of APT1, APT2, ABHD16A, and ABHD17A in male and female mouse hippocampus, striatum, cortex, and cerebellum. APT2 levels were highest in the hippocampus while APT1 and ABHD16A levels were highest and ABHD17A lowest in the cerebellum (Fig. 2A & B). This indicates brain region-specific regulation of deacylase expression, suggesting specialized roles in distinct tissues. Male and female data points largely overlapped suggesting no differential expression by sex. The only notable exception was cerebellar ABHD17A where levels trended lower in males. However, sample sizes were insufficient to analyze sex as a variable, so data were pooled. We also assessed deacylase protein levels in cultured primary rat hippocampal neurons across culture development from DIV5 to 20. All four enzymes were abundantly expressed in mature cultures, with protein levels increasing as cultures matured (Fig 2C).

**Fig. 2.**
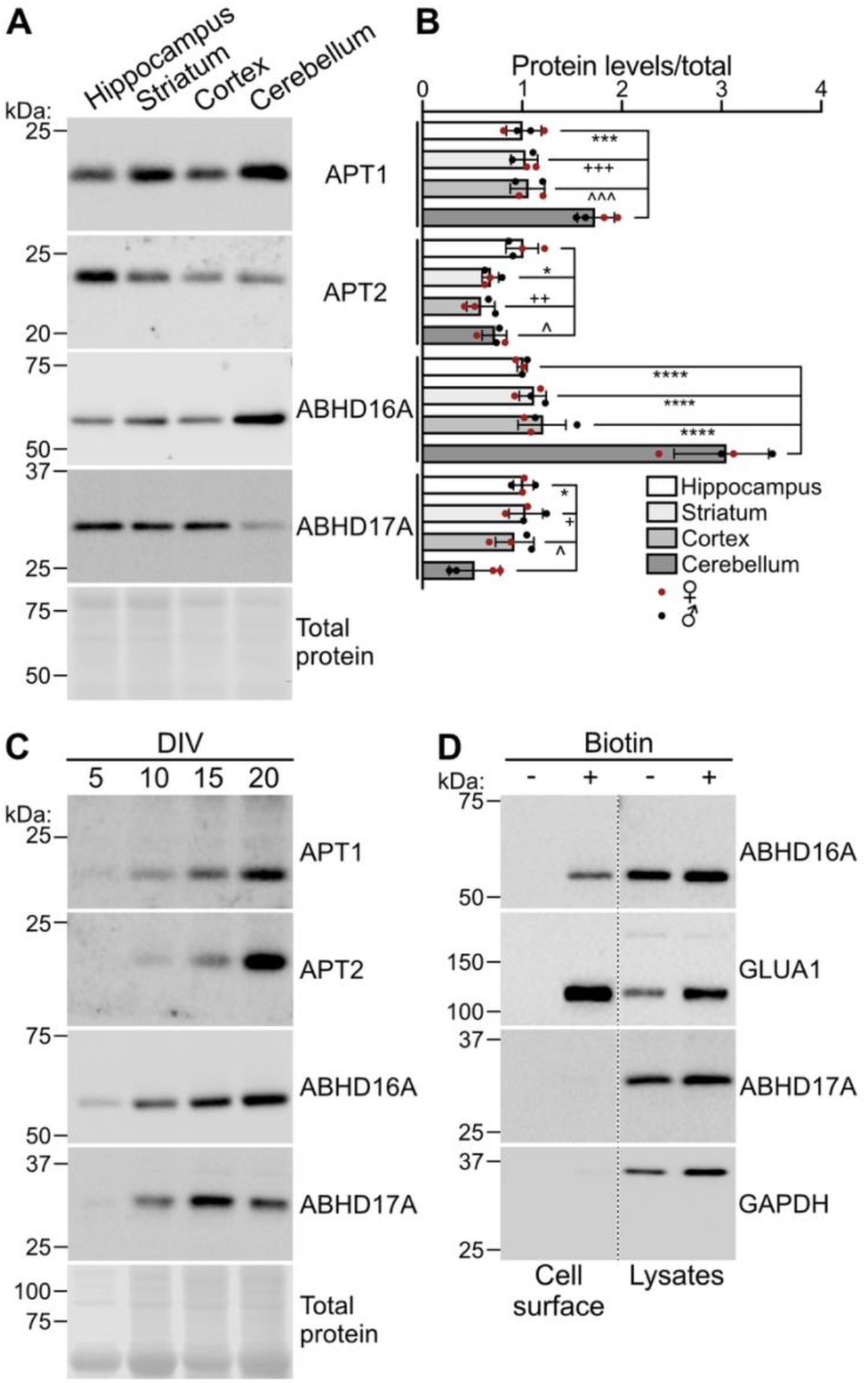
Deacylase protein levels in the mouse brain and rat hippocampal neuron culture. (**A**) APT1, APT2, ABHD16A, ABHD17A, and total (ponceau) protein levels were assessed in mouse hippocampus, striatum, cortex, and cerebellum. (**B**) Quantified data from A of indicated proteins relative to total protein normalized to hippocampus (one-way ANOVA with Tukey multiple comparisons, N=4 [2 female, 2 male]; APT1: p<0.0001, F(3,12)=19.50, *** p=0.0001, +++ p=0.0002, ^^^ p=0.0003; APT2: p=0.0042, F(3, 12)=7.58, * p=0.021, ++ p=0.0033, ^ p=0.044; ABHD16A: p<0.0001, F(3, 12)=47.90, **** p<0.0001; ABHD17A: p=0.0077, F(3,12)=6.408, * p=0.015, + p=0.011, ^ p=0.047). (**C**) APT1, APT2, ABHD16A, ABHD17A, and total (ponceau) protein levels were assessed in DIV 5, 10, 15, and 20 hippocampal neurons. (**D**) Levels of cell surface exposed protein and protein levels in parent lysates of ABHD16A and GLUA1 (positive control surface exposed protein) as well as ABHD17A and GAPDH (negative control non-surface exposed proteins) were assessed in hippocampal neurons (composite from the same images, dashed lines indicate splice points). C and D are representative of three independent experiments.

We next sought to determine the neuronal subcellular localization of the deacylases. ABHD16A is an integral membrane protein [25], so we assessed plasma membrane localization using cell surface biotinylation. ABHD16A was readily detected in the cell surface fraction with the plasma membrane glutamate receptor subunit GLUA1, whereas the cytosolic proteins ABHD17C and GAPDH were not (Fig. 2D). To broadly survey localization, hippocampal neurons were transfected to express HA-tagged deacylases expressed in the hippocampus (Fig. 1). All neurons were co-stained with Golgi-localized GM130. Because APT1 partially localizes to mitochondria in HEK293T cells [26], neurons expressing APT1-HA were additionally co-stained with mitochondrial-localized TOM20. APT1-HA displayed diffuse localization throughout the neuron with some higher-intensity puncta that overlapped minimally with either TOM20 or GM130 (Fig. 3A&B). This suggests that APT1 is largely not localized to mitochondria or the Golgi in hippocampal neurons, contrasting prior reports in non-neuronal cells [27–29]. The other deacylases were similarly detected on small puncta throughout neurons, suggesting association with various membrane compartments (Fig. 3C-H). Supporting these observations, APT1, APT2, and ABHD16A were previously identified on small motile vesicles in the brain or in cultured cortical neurons by mass spectrometry [15,30]. Further, APT1 and ABHD17A/B were previously identified on vesicles in cultured cortical and hippocampal neurons, respectively, and ABHD16A was shown to localize to the endoplasmic reticulum in cultured cerebellar neurons [11,15,25,30]. We also observed APT2-HA and HA-ABHD17C localized to the Golgi (Fig. 3B&G). For APT2, this is consistent with previous reports of Golgi localization in non-neuronal cells and cultured hippocampal neurons [11,15,27–29]. Together, these findings suggest that deacylases are broadly distributed throughout neurons, likely associating with distinct membrane microdomains and trafficking compartments, providing a foundation for future studies of endogenous neuronal deacylase localization.

**Fig 3.**
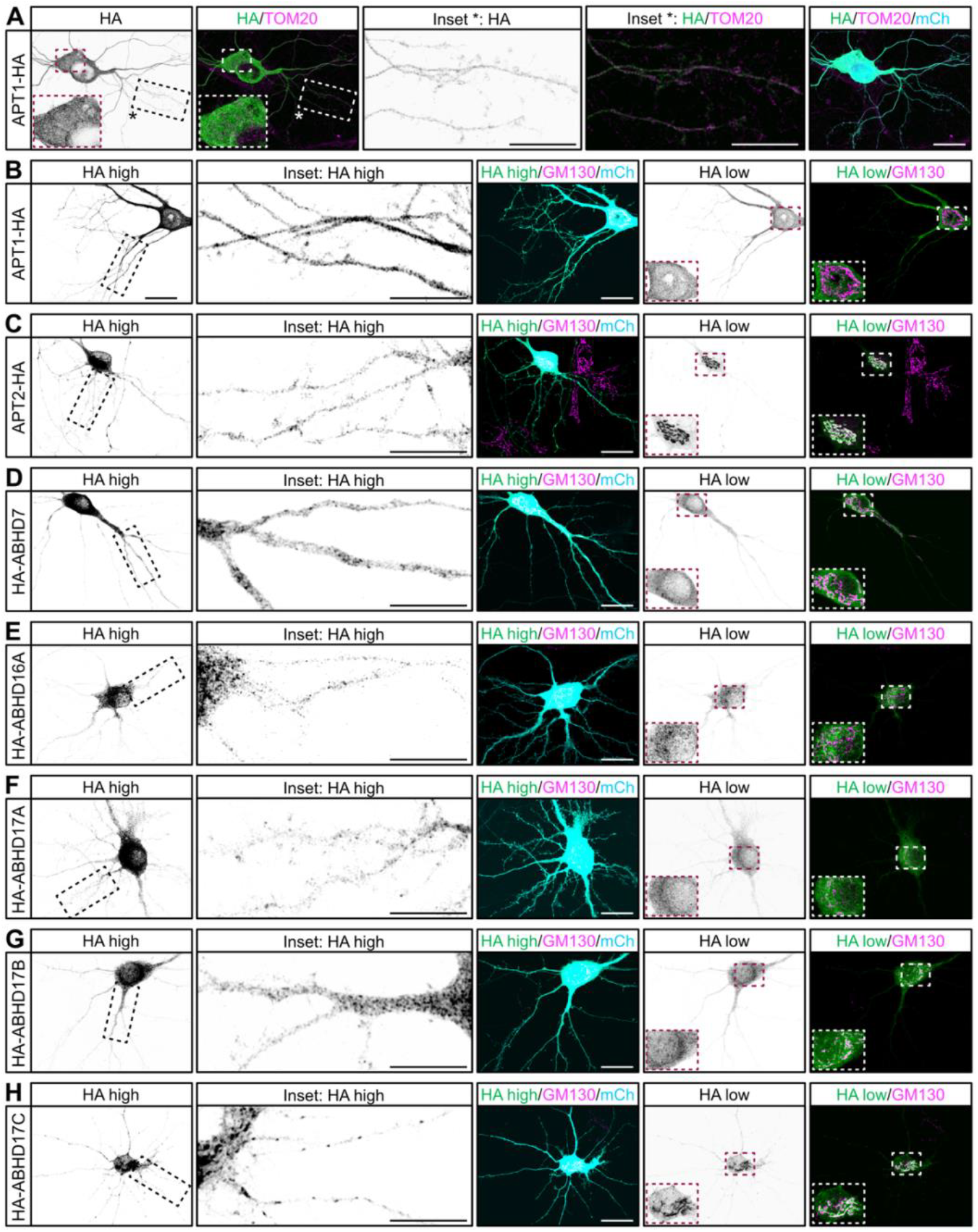
Localization of deacylases in neurons. Hippocampal neurons expressing the following HA-tagged deacylases and mCherry (mCh; cell fill) were immunostained with HA, RFP (to detect mCh), and TOM20 (mitochondria) or GM130 (Golgi) antibodies: (**A & B**) APT1-HA, (**C**) APT2-HA, (**D**) HA-ABHD7, (**E**) HA-ABHD16A, (**F**) HA-ABHD17A, (**G**) HA-ABHD17B, or (**H**) HA-ABHD17C. Black dashed boxes indicate zoomed-in regions in *‘*Inset’ panels and red and white dashed boxes indicate zoomed-in regions on the bottom left of the corresponding image. HA high and HA low are brightness-contrast adjusted versions of the same image to visualize neurite and soma structures, respectively. Scale bar in full field of view is 20 μm and in insets 10 μm.

Finally, S-acylation of deacylases has been proposed to regulate substrate access [1]. Indeed, APT1, APT2, and ABHD17B must be S-acylated to deacylate their membrane-bound substrates [11,27]. Thus, we investigated whether all ten deacylases are S-acylated using bioorthogonal labeling and click chemistry with alkyne-palmitate and alkyne-stearate in HEK293T cells. Indeed, all 10 are readily S-acylated by both fatty acids, except ABHD16A which prefers alkyne-stearate (Fig. 4). These data confirm S-acylation of the deacylases and provide a resource for selecting the optimal fatty acid in future experiments.

**Fig 4.**
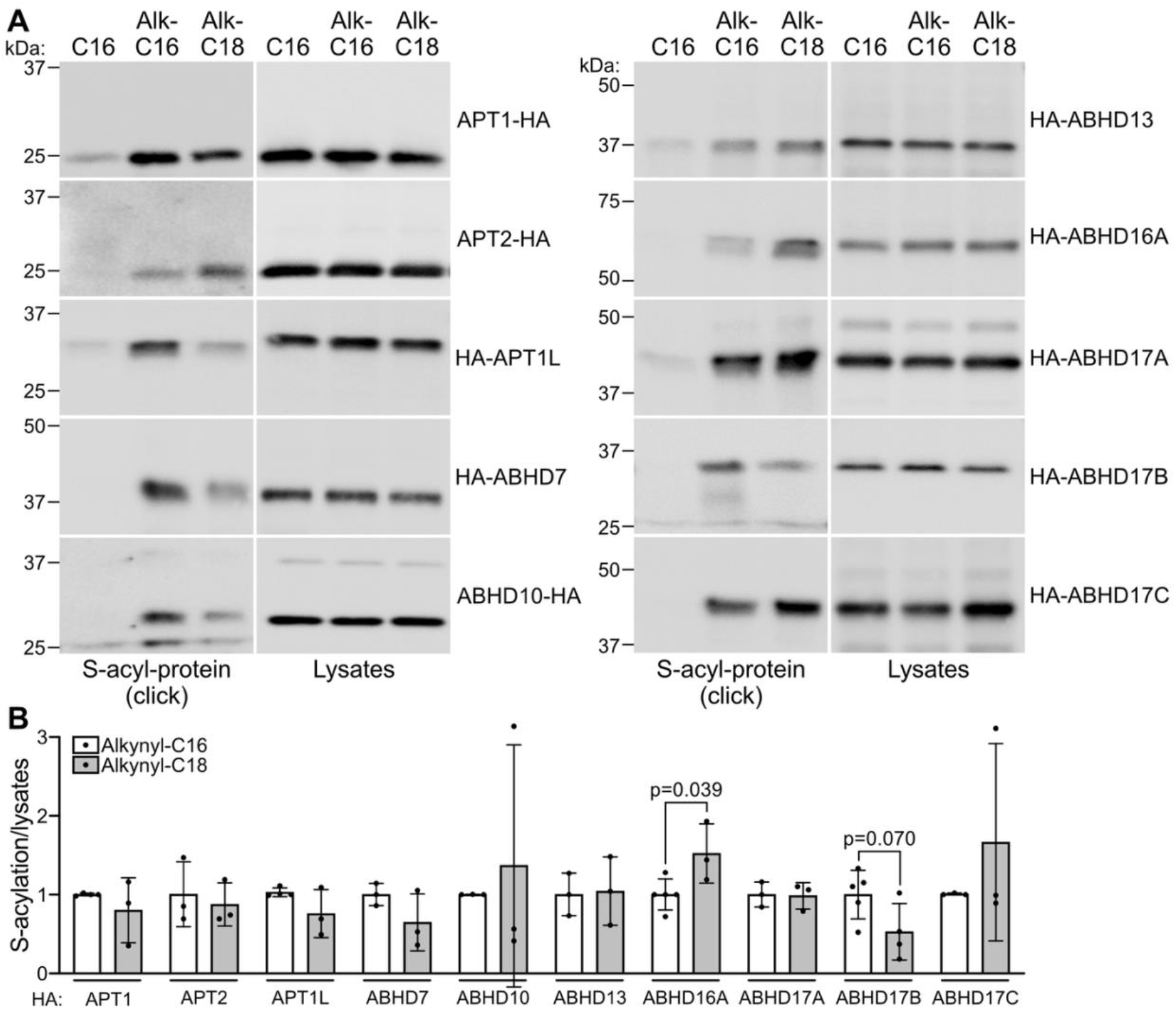
The deacylases are S-acylated in HEK293T cells. (**A**) HA-tagged deacylase-expressing HEK293T cells were labeled with palmitate (C16; negative control), alkynyl-palmitate (Alk-C16), or alkynyl-stearate (Alk-C18). S-acyl proteins were affinity purified following click chemistry. Levels of deacylase S-acylation and protein levels in parent lysates were determined. (**B**) Quantified data from A of indicated deacylase S-acylation relative to corresponding protein in parent lysates normalized to Alk-C16 (N=3-5, unpaired Student’s t test).

## Conclusions

We characterized expression and localization of deacylases across the brain and in neurons. Deacylase expression varied across brain regions, with protein levels of ABHD16A and APT1 highest in the cerebellum and APT2 in the hippocampus. Interestingly, deacylases are localized throughout neurons, likely on various membranous compartments or microdomains, and are S-acylated in HEK293T cells. Indeed, whether the deacylase neuronal localization to puncta or membrane microdomains is dependent on their S-acylation is an intriguing area for future work. Collectively, these findings provide key foundational characterizations of expression and localization of the deacylating enzymes in the brain and in neurons.

## Supplementary material

Data figures S1 and S2 as well as a key methodological resources table.

## Acknowledgments

This work was supported by the Natural Sciences and Engineering Research Council of Canada (RGPIN-2021-02547 to SSS and RGPIN-2019-04617 to DDOM). YA was supported by a Canadian Institutes of Health Research Canadian Graduate Scholarship-Masters, AHD by an Ontario Graduate Scholarship, and NNH by an Ontario Queen Elizabeth II Graduate Scholarship in Science and Technology. This research was conducted on the territory of the Mississaugas of the Credit First Nation of the Hodinöhsö:ni and Anishinaabe peoples, land that is home to First Nations, Inuit, and Métis peoples.

## CRediT

Conceptualization: GSH, NNH, AHD, CATB, SSS; Data curation: GSH, SSS; Investigation: GSH, NNH, AHD, SIMC, SSS; Formal analysis: GSH, NNH, AHD, SSS; Writing – original draft: GSH, NNH, AHD, SSS; Reviewing and editing: GSH, NNH, AHD, SIMC, JEF, YA, MR, CATB, DDOM, SSS; Resources: JEF, YA, MR; Visualization: GSH, SSS; Supervision: CATB, DDOM, SSS; Funding acquisition: DDOM, SSS

## References

[1] F.S. Mesquita, L. Abrami, M.E. Linder, S.X. Bamji, B.C. Dickinson, F.G. van der Goot, Mechanisms and functions of protein S-acylation, Nat. Rev. Mol. Cell Biol. 2209 (2024) 1–22. 10.1038/s41580-024-00700-8.

[2] A.H. Doerksen, N.N. Herath, S.S. Sanders, Fat traffic control: S-acylation in axonal transport, Mol. Pharmacol. 107 (2025) 100039. 10.1016/j.molpha.2025.100039.

[3] A.A. Petropavlovskiy, J.A. Kogut, A. Leekha, C.A. Townsend, S.S. Sanders, A sticky situation: regulation and function of protein palmitoylation with a spotlight on the axon and axon initial segment, Neuronal Signal 5 (2021) NS20210005. 10.1042/ns20210005.

[4] S.S. Sanders, D.D.O. Martin, S.L. Butland, M. Lavallée-Adam, D. Calzolari, C. Kay, J.R. Yates, M.R. Hayden, Curation of the Mammalian Palmitoylome Indicates a Pivotal Role for Palmitoylation in Diseases and Disorders of the Nervous System and Cancers, PLoS Comput. Biol. 11 (2015) e1004405–20. 10.1371/journal.pcbi.1004405.

[5] D.D.O. Martin, S.S. Sanders, Let’s get fat: emergence of S -acylation as a therapeutic target in Huntington disease, Biochem. Soc. Trans. (2024). 10.1042/bst20231290.

[6] T.A. Duarte, C.P. Ng, J.A.R. Salvador, L. Pipito, J. Greaves, V.M. Moreira, S-acylation and neuroinflammation: the therapeutic potential of zDHHC and deacylase modulation, Eur. J. Med. Chem. 303 (2026) 118429. 10.1016/j.ejmech.2025.118429.

[7] L. Mejuto, L. Pipito, C.P. Ng, N.C.O. Tomkinson, C.A. Reynolds, G. Deganutti, J. Greaves, Deacylation of SNAP25 protein family isoforms reveals distinct substrate selectivities of α/β hydrolase domain (ABHD) deacylases, bioRxiv (2026) 2026.01.21.700842. 10.64898/2026.01.21.700842.

[8] X. Shi, X. Li, Z. Xu, L. Shen, Y. Ding, S. Chen, L. Mao, W. Liu, J. Xu, ABHD16A Negatively Regulates the Palmitoylation and Antiviral Function of IFITM Proteins, Mbio (2022) e02289–22. 10.1128/mbio.02289-22.

[9] Y. Shen, L.-L. Zheng, C.-Y. Fang, Y.-Y. Xu, C. Wang, J.-T. Li, M.-Z. Lei, M. Yin, H.-J. Lu, Q.-Y. Lei, J. Qu, ABHD7-mediated depalmitoylation of lamin A promotes myoblast differentiation, Cell Rep. 43 (2024) 113720. 10.1016/j.celrep.2024.113720.

[10] D.T.S. Lin, E. Conibear, ABHD17 proteins are novel protein depalmitoylases that regulate N-Ras palmitate turnover and subcellular localization., Elife 4 (2015) e11306. 10.7554/elife.11306.

[11] N. Yokoi, Y. Fukata, A. Sekiya, T. Murakami, K. Kobayashi, M. Fukata, Identification of PSD-95 Depalmitoylating Enzymes., J. Neurosci. 36 (2016) 6431–6444. 10.1523/jneurosci.0419-16.2016.

[12] L. Tian, H. McClafferty, H.-G. Knaus, P. Ruth, M.J. Shipston, Distinct acyl protein transferases and thioesterases control surface expression of calcium-activated potassium channels., J. Biol. Chem. 287 (2012) 14718–14725. 10.1074/jbc.m111.335547.

[13] Y. Cao, T. Qiu, R.S. Kathayat, S.-A. Azizi, A.K. Thorne, D. Ahn, Y. Fukata, M. Fukata, P.A. Rice, B.C. Dickinson, ABHD10 is an S-depalmitoylase affecting redox homeostasis through peroxiredoxin-5., Nat. Chem. Biol. 15 (2019) 1232–1240. 10.1038/s41589-019-0399-y.

[14] F.L. Lemarié, N.S. Caron, S.S. Sanders, M.E. Schmidt, Y.T.N. Nguyen, S. Ko, X. Xu, M.A. Pouladi, D.D.O. Martin, M.R. Hayden, Rescue of aberrant huntingtin palmitoylation ameliorates mutant huntingtin-induced toxicity, Neurobiol Dis (2021) 105479. 10.1016/j.nbd.2021.105479.

[15] A. Virlogeux, C. Scaramuzzino, S. Lenoir, R. Carpentier, M. Louessard, A. Genoux, P. Lino, M.-V. Hinckelmann, A.L. Perrier, S. Humbert, F. Saudou, Increasing brain palmitoylation rescues behavior and neuropathology in Huntington disease mice., Science Advances 7 (2021) eabb0799. 10.1126/sciadv.abb0799.

[16] M. He, Q. Zhang, S. Chen, C. Li, B. Xie, Q. Zhao, Y. Huang, X. Fan, Expansion of the genetic and phenotypic spectrum of hereditary spastic paraplegia caused by ABHD16A gene variants: an integrated analysis based on novel variants and literature review, Front. Pediatr. 13 (2025) 1724515. 10.3389/fped.2025.1724515.

[17] D. Eshak, M. Arumugam, Unveiling therapeutic biomarkers and druggable targets in ALS: An integrative microarray analysis, molecular docking, and structural dynamic studies, Comput. Biol. Chem. 113 (2024) 108211. 10.1016/j.compbiolchem.2024.108211.

[18] E. Sjöstedt, W. Zhong, L. Fagerberg, M. Karlsson, N. Mitsios, C. Adori, P. Oksvold, F. Edfors, A. Limiszewska, F. Hikmet, J. Huang, Y. Du, L. Lin, Z. Dong, L. Yang, X. Liu, H. Jiang, X. Xu, J. Wang, H. Yang, L. Bolund, A. Mardinoglu, C. Zhang, K. von Feilitzen, C. Lindskog, F. Pontén, Y. Luo, T. Hökfelt, M. Uhlén, J. Mulder, An atlas of the protein-coding genes in the human, pig, and mouse brain, Science 367 (2020). 10.1126/science.aay5947.

[19] A.R. Wild, P. Hogg, S. Flibotte, G. Nasseri, R. Hollman, K. Haas, S.X. Bamji, BrainPalmSeq: A curated RNA-seq database of palmitoylating and de-palmitoylating enzyme expression in the mouse brain, (n.d.). 10.1101/2021.11.23.468539.

[20] G.M. Thomas, T. Hayashi, S.-L. Chiu, C.-M. Chen, R.L. Huganir, Palmitoylation by DHHC5/8 targets GRIP1 to dendritic endosomes to regulate AMPA-R trafficking., Neuron 73 (2012) 482–496. 10.1016/j.neuron.2011.11.021.

[21] T.R. Suk, C.E. Part, J.L. Zhang, T.T. Nguyen, M.M. Heer, A. Caballero-Gómez, V.S. Grybas, P.M. McKeever, B. Nguyen, T. Ali, S.M. Callaghan, J.M. Woulfe, J. Robertson, M.W.C. Rousseaux, A stress-dependent TDP-43 SUMOylation program preserves neuronal function, Mol. Neurodegener. 20 (2025) 38. 10.1186/s13024-025-00826-z.

[22] E.K. Brinkman, T. Chen, M. Amendola, B. van Steensel, Easy quantitative assessment of genome editing by sequence trace decomposition, Nucleic Acids Res. 42 (2014) e168–e168. 10.1093/nar/gku936.

[23] J. Schindelin, I. Arganda-Carreras, E. Frise, V. Kaynig, M. Longair, T. Pietzsch, S. Preibisch, C. Rueden, S. Saalfeld, B. Schmid, J.-Y. Tinevez, D.J. White, V. Hartenstein, K. Eliceiri, P. Tomancak, A. Cardona, Fiji: an open-source platform for biological-image analysis., Nat Meth 9 (2012) 676–682. 10.1038/nmeth.2019.

[24] C.A. Townsend, A.A. Petropavlovskiy, J.A. Kogut, A.M. Church, S.S. Sanders, Protocol to identify S-acylated proteins in hippocampal neurons using ω-alkynyl fatty acid analogs and click chemistry, STAR Protoc. 5 (2024) 103068. 10.1016/j.xpro.2024.103068.

[25] S. Singh, A. Joshi, S.S. Kamat, Mapping the Neuroanatomy of ABHD16A, ABHD12, and Lysophosphatidylserines Provides New Insights into the Pathophysiology of the Human Neurological Disorder PHARC, Biochemistry 59 (2020) 2299–2311. 10.1021/acs.biochem.0c00349.

[26] R.S. Kathayat, Y. Cao, P.D. Elvira, P.A. Sandoz, M.-E. Zaballa, M.Z. Springer, L.E. Drake, K.F. Macleod, F.G.V.D. Goot, B.C. Dickinson, Active and dynamic mitochondrial S-depalmitoylation revealed by targeted fluorescent probes., Nature Communications 9 (2018) 334. 10.1038/s41467-017-02655-1.

[27] L. Abrami, M. Audagnotto, S. Ho, M.J. Marcaida, F.S. Mesquita, M.U. Anwar, P.A. Sandoz, G. Fonti, F. Pojer, M.D. Peraro, F.G.V.D. Goot, Palmitoylated acyl protein thioesterase APT2 deforms membranes to extract substrate acyl chains, Nat. Chem. Biol. (2021) 1–27. 10.1038/s41589-021-00753-2.

[28] N. Vartak, B. Papke, H.E. Grecco, L. Rossmannek, H. Waldmann, C. Hedberg, P.I.H. Bastiaens, The Autodepalmitoylating Activity of APT Maintains the Spatial Organization of Palmitoylated Membrane Proteins, Biophysj 106 (2014) 93–105. 10.1016/j.bpj.2013.11.024.

[29] E. Kong, S. Peng, G. Chandra, C. Sarkar, Z. Zhang, M.B. Bagh, A.B. Mukherjee, Dynamic palmitoylation links cytosol-membrane shuttling of acyl-protein thioesterase-1 and acyl-protein thioesterase-2 with that of proto-oncogene H-Ras product and growth associated protein-43., J. Biol. Chem. (2013). 10.1074/jbc.m112.421073.

[30] M.-V. Hinckelmann, A. Virlogeux, C. Niehage, C. Poujol, D. Choquet, B. Hoflack, D. Zala, F. Saudou, Self-propelling vesicles define glycolysis as the minimal energy machinery for neuronal transport., Nature Communications 7 (2016) 13233. 10.1038/ncomms13233.

