## Supplementary material for "Brain and neuronal expression and localization of de-S-acylating enzymes"

**Table 1.** Key Resources table.

**DNA**

| **Plasmid** | **Source** | **Species and promoter** |
|---|---|---|
| pEF-BOS-HA-*LYPLA1* | Dr. Jennifer Greaves, Coventry University [1] | Human cDNA; EF1α (human eukaryotic elongation factor 1α) |
| pEF-BOS-HA-*LYPLA2* | Dr. Jennifer Greaves [1] | Human cDNA; EF1α |
| pEF-BOS-HA-*LYPLA1L* | Dr. Jennifer Greaves [1] | Human cDNA; EF1α |
| pEF-BOS-HA-EPHX4 | cDNA from DNASU, #HsCD00289175 [2] | Human cDNA; EF1α |
| pEF-BOS-HA-*ABHD10* | Dr. Jennifer Greaves [1] | Human cDNA; EF1α |
| pEF-BOS-HA-*ABHD13* | Dr. Jennifer Greaves [1] | Human cDNA; EF1α |
| pEF-BOS-HA-*Abhd16A* | Dr. Jennifer Greaves [3] | Mouse cDNA; EF1α |
| pEF-BOS-HA-*ABHD17A* | Dr. Jennifer Greaves [1] | Human cDNA; EF1α |
| pEF-BOS-HA-*ABHD17B* | cDNA from DNASU, #HsCD00861137 [2] | Human cDNA; EF1α |
| pEF-BOS-HA-*ABHD17C* | Dr. Jennifer Greaves [1] | Mouse cDNA; EF1α |
| pFEW-HA | Dr. Gareth Thomas, Temple University [4] | EF1α |
| pFEW-*LYPLA1*-HA | This paper | Human cDNA; EF1α |
| pFEW-*LYPLA2*-HA | This paper | Human cDNA; EF1α |
| pFEW-*ABHD10*-HA | This paper | Human cDNA; EF1α |
| pFEmCherryW | Dr. Gareth Thomas [5] | mCherry fluorescent protein, EF1α |
| LentiCas9-Blast | Dr. Feng Zhang (Addgene plasmid #52962) [6] | *Streptococcus pyogenes*; EF1α |
| pCLIP-DUAL-SFFV-ZsGreen-sgRNA | University of Ottawa Genome Editing and Molecular Biology (GEM) Facility [7] | Human gene targeting gRNAs; U6 RNA polymerase III |

**gRNA sequences**

| **Gene** | **Sequences** |
|---|---|
| *LYPLA1* | gRNAa GGGCACGATGGCGGGCAGCG and gRNAb CCAAAGGGTCACAATCCCCG |
| *LYPLA1* | gRNAa GGCCATACTCCAGTGCCATG and gRNAb AACGCCACGATGCCAGCCAG |
| *ABHD16A* | gRNAa GCATCTCAATCTAAACAACG and gRNAb CAGGGGCTCTGGGCGAAGCA |
| *eGFP* | Non-targeting negative control, gRNAa CGAGGAGCTGTTCACCGGGG and gRNAb CAACTACAAGACCCGCGCCG |

**TIDE primers**

| **Name** | **Sequence** |
|---|---|
| *LYPLA1*_gRNAa_F3 | TGTGAGCTGAGGCGGTGTATGT |
| *LYPLA1*_gRNAa_R3 | AGGCGATTTCCTGGCGGACTTA |
| *LYPLA1*_gRNAb_F | TTTTGCAGGGTCCTATCGGT |
| *LYPLA1*_gRNAb_R | ACACCTACCTGTTGACACGA |
| *LYPLA2*_gRNAa_F | CTAGCTTTGCCCTGAGTCCT |
| *LYPLA2*_gRNAa_R | AAGTGGGAGTGGGAAATGGT |
| *LYPLA2*_gRNAb_F | TCATGACCCCTCTCTCTCCT |
| *LYPLA2*_gRNAb_R | CGAGGATGCAGACACAAAGG |
| *ABHD16A*_gRNAa_F | TCTGCCATCCCTTCACCTTT |
| *ABHD16A*_gRNAa_R | GGCCTTTTACCAACTTGCAC |
| *ABHD16A*_gRNAb_F | CGTTCCTAGACCCCACTGTT |
| *ABHD16A*_gRNAb_R | CAAGGCAGCTTCTTAACCCG |

| **Antibodies**<br>(IB=immunoblot; ICC=immunocytochemistry) | | |
|---|---|---|
| **Target, type, & dilution** | **Catalogue number** | **Manufacturer** |
| Anti-HA rabbit monoclonal; ICC 1:500; IB 1:5000 | #S3724 | Cell Signaling Technologies, Danvers, MA, USA |
| Anti-HA mouse monoclonal; ICC 1:500; IB 1:1000 | HA.11, #901514 | BioLegend, San Diego, CA, USA |
| Anti-BAT5/ABHD6A rabbit monoclonal; ICC 1:500 IB 1:1000 | #Ab185549 | Abcam, Cambridge, UK |
| Anti-GM130 mouse monoclonal; ICC 1:500 | #610822 | Waters Biosciences, Milford, MA, USA |
| Anti-RFP (red fluorescent protein) rat monoclonal; ICC 1:1000 | #5f8 | Proteintech, Rosemond, IL, USA |
| Anti-TOM20 rabbit polyclonal; ICC 1:200 | #11802-1-AP | Proteintech |
| Anti-APT1 rabbit polyclonal; IB 1:1000 | #16055-1 | Proteintech |
| Anti-LYPLA1 rabbit monoclonal; IB 1:1000 | #Ab91606 | Abcam |
| Anti-APT2 rabbit polyclonal; IB 1:1000 | #PA5-27653 | ThermoFisher Scientific, Waltham, MA, USA |
| Anti-ABHD7 rabbit polyclonal; IB 1:1000 | #PA5-106586 | ThermoFisher Scientific |
| Anti-ABHD10 rabbit polyclonal; IB 1:1000 | #STJ190492 | St. John's Laboratory, London, UK |
| Anti-ABHD13 rabbit polyclonal; IB 1:1000 | #STJ190493 | St. John's Laboratory |
| Anti-ABHD13 rabbit polyclonal; IB 1:1000 | #PA5-75745 | ThermoFisher Scientific |
| Anti-ABHD17A rabbit polyclonal; IB 1:1000 | #15854-1-AP | Proteintech |
| Anti-ABHD17A rabbit polyclonal; IB 1:1000 | #STJA0007854 | St. John's Laboratory |
| Anti-ABHD17C rabbit polyclonal; IB 1:1000 | #STJ198183 | St. John's Laboratory |
| Alexa Fluor 568-conjugated goat anti-rat; ICC 1:500 | #A-11077 | ThermoFisher Scientific |
| Alexa Fluor 488-conjugated goat anti-rabbit; ICC 1:500 | #A-11008 | ThermoFisher Scientific |
| Alexa Fluor 647-conjugated goat anti-mouse IgG1; ICC 1:500 | #A-21240 | ThermoFisher Scientific |
| Alexa Fluor 488-conjugated goat anti-mouse IgG1; ICC 1:500 | #A-21121 | ThermoFisher Scientific |
| Alexa Fluor 647-conjugated goat anti-rabbit; ICC 1:500 | #A-21244 | ThermoFisher Scientific |
| HRP (horseradish peroxidase)-conjugated donkey anti-rabbit; IB 1:5000 | #AP182P | MilliporeSigma |
| HRP-conjugated horse anti-mouse; IB 1:5000 | #7076 | Cell Signaling Technologies |
| Alexa Fluor 680-conjugated goat anti-mouse; IB 1:5000 | #A21057 | ThermoFisher Scientific |
| **Media, reagents, & chemicals** | | |
| **Reagent** | **Catalogue number** | **Manufacturer** |
| Neurobasal media | #21103049 | ThermoFisher Scientific |
| B-27 Supplement | #17504044 | ThermoFisher Scientific |
| 5-Fluoro-2'-deoxyuridine | #2F0503 | MilliporeSigma, Burlington, MA, USA |
| Poly-L-lysine hydrobromide | #P2636 | MilliporeSigma |

| | | |
|---|---|---|
| EZ-Link Sulfo-NHS-SS-Biotin | #21331 | ThermoFisher Scientific |
| DMEM (Dulbecco's modified eagle medium) | #319-015-CL | Wisent Inc., Saint-Jean-Baptiste, QC, Canada |
| IMDM (Iscove's Modified Dulbecco's medium) | #319-105-CL | Wisent Inc. |
| Penicillin-Streptomycin | #15140122 | ThermoFisher Scientific |
| Glutamax Supplement | #35050061 | ThermoFisher Scientific |
| L-glutamine | #609-065-EL | Wisent Inc. |
| Lipofectamine 2000 Transfection reagent | #11668027 | ThermoFisher Scientific |
| PEI | #919012 | MilliporeSigma |
| 32% paraformaldehyde | #15714S | Electron Microscopy Sciences, Morgantown, PA, USA |
| PermaFluor Aqueous Mounting Medium | #TA-006-FM | Epredia, Kalamazoo, MI, USA |
| Puromycin | #PUR333.25 | BioShop Canada Inc, Burlington, ON, Canada |
| Fetal bovine serum (FBS) | #12483020 | Gibco, Waltham, MA, USA |
| Charcoal stripped FBS | #F6765 | MilliporeSigma, Burlington, MA, USA |
| Alkynyl-palmitate (15-hexadecynoic acid) | #CCT-1165 | Vector Laboratories, Newark, CA, USA |
| Alkynyl-stearate (17-octadecynoic acid) | #CCT-1166 | Vector Laboratories |
| Palmitate | #P5585 | MilliporeSigma |
| Fatty acid free BSA | #A7030 | MilliporeSigma |
| Roche cOmplete EDTA-free protease inhibitor cocktail | #11873580001 | MilliporeSigma |
| *DC* (detergent compatible) protein assay | #5000111 | Bio-Rad Laboratories Inc, Hercules, CA, USA |
| Biotin azide plus | #CCT-1488 | Vector Laboratories |
| Q5 High-Fidelity DNA Polymerase | #M0491L | New England Biolabs (NEB), Ipswich, MA, USA |
| Monarch® Spin PCR & DNA Cleanup Kit | #T1130S | NEB |
| Proteinase K, Molecular Biology Grade | #P8107S | NEB |
| TBTA | #678937 | MilliporeSigma |
| $CuSO_4$ | #209198 | MilliporeSigma |
| Sodium L-ascorbate | #7631 | MilliporeSigma |
| Leupeptin | #LEU001 | BioShop Canada Inc., Burlington, ON, Canada |
| Benzamidine | #BEN666 | BioShop Canada Inc. |
| Pierce high capacity NeutrAvidin agarose beads | #PI29204 | ThermoFisher Scientific |
| Hydroxylamine hydrochloride | #159417 | MilliporeSigma |
| 0.45 µm Cytiva Amersham Protran NC Nitrocellulose Membrane | #10600002 | Cytiva, Marlborough, MA, USA |
| Clarity Western ECL substrate | #1705061 | Bio-Rad |
| **Buffers & media composition** | | |
| **Buffer/media** | **Composition** | |
| Homogenization Buffer | 10 mM phosphate buffer pH 7.4, 0.32 M sucrose, 1 mM EDTA (ethylenediaminetetraacetic acid), 6 M Urea | |
| 2% SDS buffer (2SB) | 2% SDS, 50 mM pH 7.0 HEPES (N-2-hydroxyethylpiperazine-N′-2-ethanesulfonic acid), 1 mM EDTA | |
| 2x HBS | 270 mM NaCl, 1.5 mM $Na_2HPO_4•7H_2O$, 40 mM HEPES, 10 mM KCl, 10 mM D-glucose, pH 7.0 | |
| Parafix | 4% paraformaldehyde, 4% sucrose, 1x PBS | |
| Artificial cerebral spinal fluid (aCSF) | 25 mM HEPES pH 7.4, 120 mM NaCl, 5 mM KCl, 2 mM $CaCl_2$, 20 mM glucose, 1 mM $MgCl_2$ | |
| Dilution buffer | 50 mM HEPES pH 7.0, 1% Triton X-100, 1 mM EDTA, 1 mM | |

| | |
|---|---|
| | EGTA ethylene glycol-bis(β-aminoethyl ether)-N,N,N′,N′-tetraacetic acid], 1 μg/mL leupeptin and 1 mM benzamidine |
| Elution buffer | Dilution buffer with 0.2% SDS, 250 mM NaCl, and 1% β-mercaptoethanol |
| Hydroxylamine elution buffer | 1M $NH_2OH$, 50mM HEPES pH 7.4, 150mM NaCl, 0.1% SDS |
| Genomic DNA buffer | 100 mM Tris pH 8.0, 50 mM EDTA, 40 mM NaCl, 0.2% SDS |
| Complete Neurobasal Media | Neurobasal medium with 1% GlutaMAX Supplement, 2% B-27 Supplement, and 1% penicillin-streptomycin (P/S) |
| Complete DMEM | DMEM, 10% fetal bovine serum [FBS], 1% P/S, 1% L-glutamine |
| Complete IMDM | IMDM, 10% FBS, 1% P/S, 1% L-glutamine |
| **Software & webtools** | |
| TIDE (Tracking of Indels by Decomposition) online tool | https://tide.nki.nl/ |
| Fiji | ImageJ |
| Image Studio Software version 6 | Li-COR Biotech, Lincoln, NE, USA |
| Image Lab software | Bio-Rad |
| Prism 11 | GraphPad Software, Bosten, MA, USA |

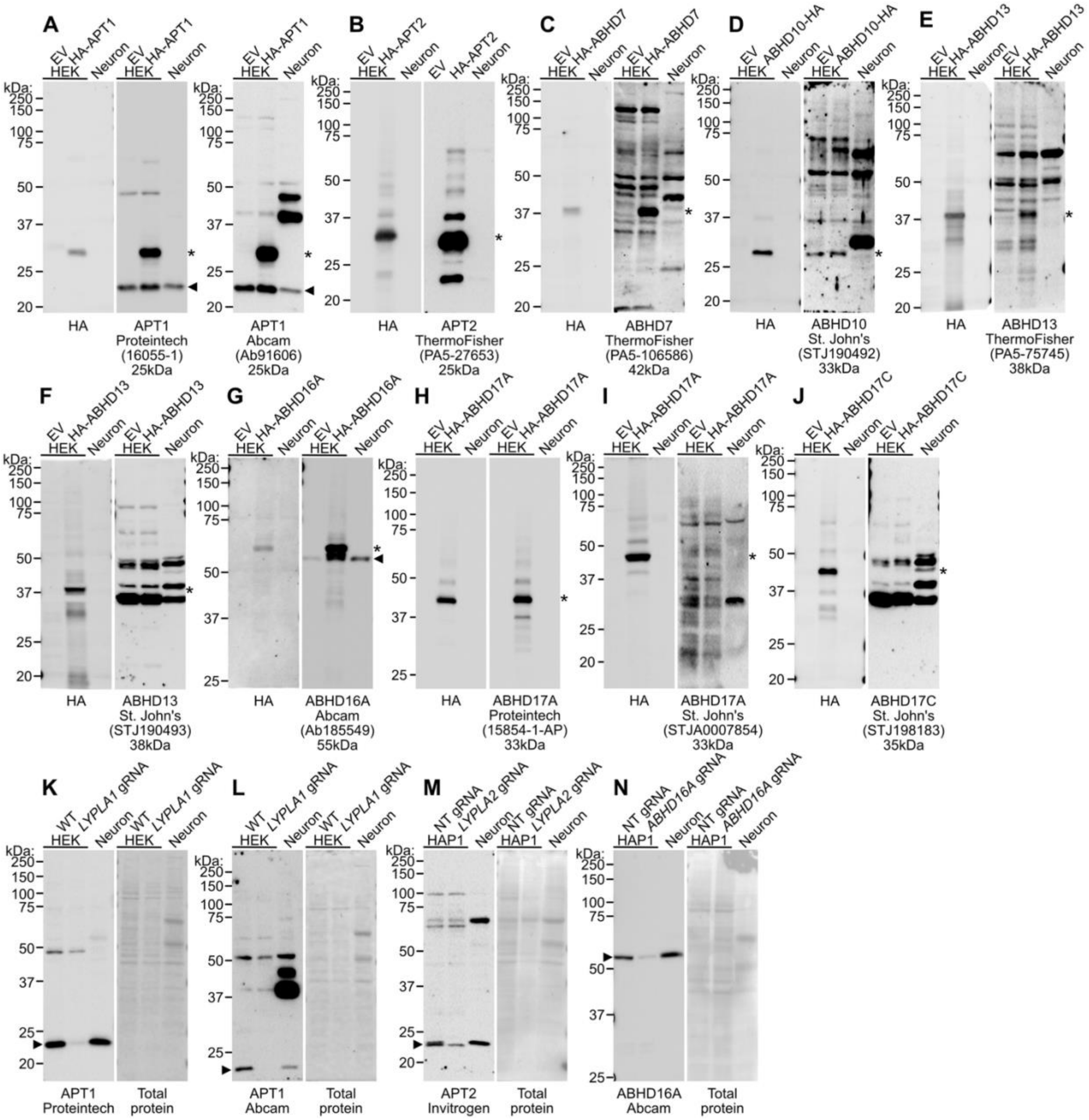


**Fig. S1.** Deacylase antibody testing and validation for immunoblot. (**A-I**) Lysates from HEK293T cells transfected with empty vector (EV) or the indicated HA-tagged deacylases as well as from hippocampal neuron cultures were subjected to SDS-PAGE and immunoblotting using the following antibodies: HA (left panels in A-I), (**A**) Proteintech APT1 (middle panel), Abcam APT1 (right panel), (**B**) Invitrogen APT2 (right panel), (**C**) Invitrogen ABHD7 (right panel), (**D**) St. Johns ABHD10 (right panel), (**E**) Invitrogen ABHD13 (right panel), (**F**) St. Johns ABHD13 (right panel), (**G**) Abcam ABHD16A (right panel), (**H**) Proteintech ABHD17A (right panel), (**I**) St. Johns ABHD17A (right panel), or (**J**) St. Johns ABHD17C (right panel). The catalogue numbers for each antibody and the predicted molecular weight are indicated below each blot image, the Asterix indicates the HA-tagged protein band, and the arrowhead indicates the endogenous protein band if visible (for APT2 and ABHD17A antibody binding to the overexpressed HA-tagged protein masks binding to the endogenous protein). (K-L) Lysates from wild type and *LYPLA1* knockout (KO) polyclonal HEK293T cells were subjected to SDS-PAGE and immunoblot using the following antibodies: (**K**) Proteintech APT1 (left panel) or (**L**) Abcam APT1 (left panel). (**M**) Lysates from NT (non-targeting gRNA) and *LYPLA2* KO polyclonal human haploid (HAP1) cells were subjected to SDS-PAGE and immunoblot using an antibody against APT2 (left panel). (**N**) Lysates from NT (non-targeting gRNA) and *ABHD16A* KO polyclonal HAP1 cells were subjected to SDS-PAGE and immunoblot using an antibody against ABHD16A (left panel). Total protein in K-N was ponceau stain (right panels) and the arrowhead indicates the endogenous protein band.

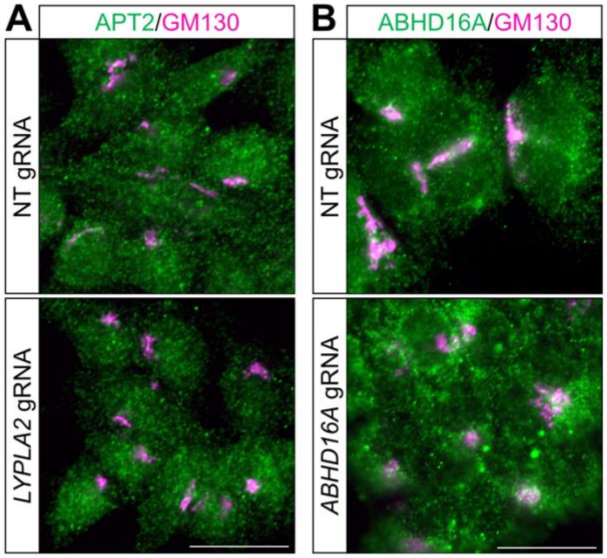


**Fig. S2.** Deacylase antibody testing and validation for immunocytochemistry. (**A**) NT (top panel) and *LYPLA2* KO (bottom panel) HAP1 cells were fixed and immunostained with APT2 (green) and GM130 (Golgi; magenta) antibodies. (**B**) NT (top panel) and *ABHD16A* KO (bottom panel) HAP1 cells were fixed and immunostained with ABHD16A (green) and GM130 (magenta) antibodies. Scale bar is 20 µm.

**Table S1.** Indel frequency analysis of knockout polyclonal cell lines. Purified PCR amplicons including each gRNA targeted sequence from NT and the indicated KO HEK293T or HAP1 cells were Sanger sequenced and the resulting chromatograms analyzed using the TIDE online tool.

| **Gene** | **gRNAa** | **gRNAb** |
|---|---|---|
| *LYPLA1* | 81% | 71% |
| *LYPLA2* | 89% | 97% |
| *ABHD16A* | 50% | 13% |

**References**

[1] L. Mejuto, L. Pipito, C.P. Ng, N.C.O. Tomkinson, C.A. Reynolds, G. Deganutti, J. Greaves, Deacylation of SNAP25 protein family isoforms reveals distinct substrate selectivities of α/β hydrolase domain (ABHD) deacylases, bioRxiv (2026) 2026.01.21.700842. https://doi.org/10.64898/2026.01.21.700842.

[2] C.Y. Seiler, J.G. Park, A. Sharma, P. Hunter, P. Surapaneni, C. Sedillo, J. Field, R. Algar, A. Price, J. Steel, A. Throop, M. Fiacco, J. LaBaer, DNASU plasmid and PSI:Biology-Materials repositories: resources to accelerate biological research, Nucleic Acids Res. 42 (2014) D1253–D1260. https://doi.org/10.1093/nar/gkt1060.

[3] T.J. Ahonen, C.P. Ng, B. Farinha, B. Almeida, B.L. Victor, C. Reynolds, E. Kalso, J. Yli-Kauhaluoma, J. Greaves, V.M. Moreira, Probing the Interactions of Thiazole Abietane Inhibitors with the Human Serine Hydrolases ABHD16A and ABHD12, ACS Med. Chem. Lett. 14 (2023) 1404–1410. https://doi.org/10.1021/acsmedchemlett.3c00313.

[4] S.M. Holland, K.M. Collura, A. Ketschek, K. Noma, T.A. Ferguson, Y. Jin, G. Gallo, G.M. Thomas, Palmitoylation controls DLK localization, interactions and activity to ensure effective axonal injury signaling., Proceedings of the National Academy of Sciences 113 (2016) 763–768. https://doi.org/10.1073/pnas.1514123113.

[5] K.M. Collura, J. Niu, S.S. Sanders, A. Montersino, S.M. Holland, G.M. Thomas, The palmitoyl acyltransferases ZDHHC5 and ZDHHC8 are uniquely present in DRG axons and control retrograde signaling via the Gp130/JAK/STAT3 pathway., Journal of Biological Chemistry (2020) jbc.RA120.013815. https://doi.org/10.1074/jbc.ra120.013815.

[6] N.E. Sanjana, O. Shalem, F. Zhang, Improved vectors and genome-wide libraries for CRISPR screening, Nat. Methods 11 (2014) 783–784. https://doi.org/10.1038/nmeth.3047.

[7] N. Erard, S.R.V. Knott, G.J. Hannon, A CRISPR Resource for Individual, Combinatorial, or Multiplexed Gene Knockout, Mol. Cell 67 (2017) 348-354.e4. https://doi.org/10.1016/j.molcel.2017.06.030.
